# Pluripotent stem cell-derived NK progenitor cell therapy prevents tumour occurrence and eradicates minimal residual disease

**DOI:** 10.1101/2025.01.07.631650

**Authors:** Zhiqian Wang, Yunqing Lin, Dehao Huang, Leqiang Zhang, Chengxiang Xia, Qitong Weng, Yanhong Liu, Tongjie Wang, Mengyun Zhang, Jiaxin Wu, Hanmeng Qi, Lijuan Liu, Yiyuan Shen, Yi Chen, Yanping Zhu, Fangxiao Hu, Jinyong Wang

## Abstract

The lack of persistence and short-term efficacy presents a major challenge for CAR-NK cell therapy. Here, we addressed this issue by developing pluripotent stem cell-derived NK lineage-committed progenitor (iNKP) cell therapy. For the first time, we generated abundant iNKP cells via an organoid culture system. The iNKP cells, engineered to express CXCR4 and chimeric antigen receptors (CAR), efficiently migrated to the bone marrow and generated CAR-iNK cells persisting in peripheral blood (PB) for over 80 days. Notably, CAR-iNKP cell therapy durably protected animals from tumour occurrence. Furthermore, a single low-dose infusion of CAR-iNKP cells following conventional chemotherapy eradicated minimal residual disease (MRD), leading to long-term complete remission. Our findings present a novel strategy to overcome the limitations of traditional CAR-NK cell therapy and offer dural breakthroughs for the prevention of tumour occurrence and relapse.

## Introduction

Immunotherapy has demonstrated significant clinical potential in treating tumours and autoimmune diseases^1,2^. Natural killer (NK) cells belong to the innate immune system and can inherently kill a broad spectrum of tumour and virus-infected cells^3^. NK cells can also kill target cells through antibody-dependent cell-mediated cytotoxicity (ADCC) or CAR-mediated cytotoxicity^4,5^. Patients receiving autologous or allogeneic NK cells do not experience severe side effects such as cytokine release syndrome or neurotoxicity^3^. NK and CAR-NK cells have already been widely used in clinical trials to treat cancer patients^6^. Therefore, NK cells possess unique features complementing T cells in immune cell therapy.

One of the significant challenges in the treatment of malignant tumours is tumour relapse. MRD after conventional radiation, chemotherapy, and CAR-T cell therapies are the primary causes of tumour recurrence^7^. Studies have shown a significant inverse correlation between NK cell activity and tumour incidence^8^. Moreover, NK cell dysfunction is a critical factor in tumour recurrence and metastasis, and NK cell infusion following radiation or chemotherapy can effectively reduce recurrence rates in patients^9^. Thus, NK cells have the potential for tumour prevention and the elimination of MRD.

Autologous-infused immune-compatible NK cells persist in PB for about two weeks^10^. Conventional NK cell therapy requires high doses and repeated infusions to enhance treatment outcomes. The infusion doses of NK cells are as high as 10^7^ to 10^9^ cells per kilogram^11^. In clinical trials for CD19 CAR-NK cells targeting B-cell malignancies (NCT05020678), a single infusion dose reaches 1.5 billion cells^12^. The requirement for high doses and multiple infusions directly adds to the cost and complexity of NK cell therapy, ultimately affecting its long-term efficacy, affordability, and accessibility.

This study develops a new pluripotent stem cell-derived iNKP cell therapy. iNKP cell infusion overcomes the short-term persistence problem faced by injecting mature NK cells. CXCR4-expressing iNKP (R4-iNKP) cells have a superior capability to migrate to bone marrow for development. A very low single dose (2 × 10^5^ iNKP cells/dose) of iNKP cell infusion resulted in abundant iNK cells in the recipient’s PB over 80 days before declining to immeasurable levels. Furthermore, by introducing CAR expression elements at the pluripotent stem cell stage, the derived CAR-R4iNKP cells engrafted and persistently produced high levels of CAR-iNK cells in PB and multiple organs *in vivo*. Further, a single dose of CAR-R4iNKP cell infusion prevented tumour occurrence following tumour injection challenges. Combined with traditional chemotherapy, a single dose of CAR-R4iNKP cell infusion eradicated the MRD, resulting in long-term remission of tumour-bearing animals. Pluripotent stem cell-derived iNKP therapy has the prospect of solving the short-term efficacy of infusing mature NK cells, significantly improving precision prevention of cancers, and eradicating MRD.

## Results

### Pluripotent stem cell-derived iNKP cells engraft and generate NK cells *in vivo*

Infused NK cells persist shortly in the body, ranging from a few days to two weeks, and lack sustained efficacy in treating tumours. Although high doses and multiple doses regimens were applied to improve therapeutic efficacy, the inherent fast-replenishing rate of NK cells remains a great challenge for prolonged treatment outcomes. We propose that infusion of NK-lineage-restricted progenitor cells might give rise to a prolonged wave of NK lymphopoiesis *in vivo* and overcome the persistency problem faced by infusion of mature NK cells. To test our hypothesis, we first attempted to identify the induced NK-lineage committed progenitor cells from pluripotent stem cells using an organoid induction system we recently established for the large-scale generation of iNK cells^13^. We isolated the single cells from the induction organoids daily from day 16 to day 20 and performed single-cell RNA sequencing (**Fig. 1a**). After quality control, 54,624 cells were retained for downstream data analysis. Six cell populations were identified based on cell-type classical markers, including induced stromal cells (iStroma) (PDGFRA)^14^, induced endothelial cells (iEndo) (CDH5)^15,16^, induced hematopoietic progenitor cells (iHPC) (ANGPT1, SPINK2)^17–19^, induced megakaryocytic–erythroid cells (iMkE) (GATA1, GATA2)^20^, induced myeloid cells (iMyeloid) (SPI1)^21^, and iNK lineage cells (CD34, CD38, CD7, NKG7, IL2RB, and GNLY)^22–24^ ((Extended Data Fig. 1a,b). To further resolve iNKP subpopulations, we extracted the subsets of cells co-expressing *CD45* and *CD34* and those with NK lineage features. We further integrated our data with publicly available NKP (164 cells) and NK cell (166 cells) transcriptomic datasets (GSE149938) ^25^. UMAP and pseudo-time analysis defined early-iNKP (e-iNKP), late-iNKP (l-iNKP), and immature-iNK cell (im-iNK) subsets. The remaining *CD34-*positive progenitors, which included populations with myeloid and erythroid-megakaryocytic progenitor features, were classified as other-iHPC. e-iNKP appeared on day 16 (23.7%) and peaked on day 17 (28.1%). l-iNKP and im-iNK populations emerged on day 17 (1.9% and 1.5%) and peaked on day 20 (20.9% and 52.8%). By day 19, the total NK lineage-restricted progenitors (e-iNKP and l-iNKP) peaked (35.4%). Pseudotime analysis indicated that NK cell development followed a trajectory of e-iNKP, l-iNKP, and im-iNK (**Fig. 1b,c and Supplementary table 1**).

**Fig. 1.**
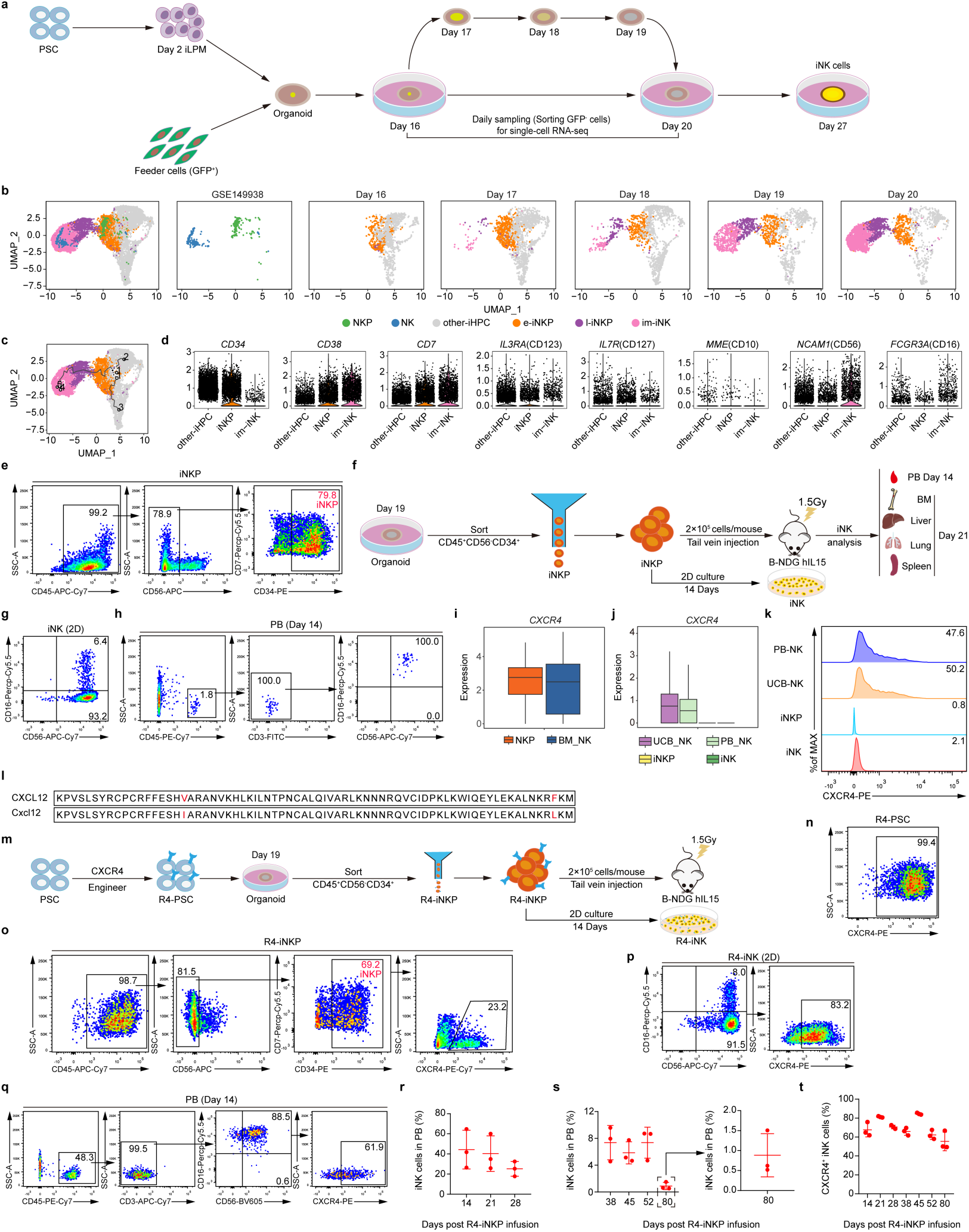
iNKP cells expressing CXCR4 efficiently engrafted and generated mature iNK cells over 80 days *in vivo*. **a,** Schematic diagram of single-cell RNA-seq assays for dissecting NK progenitor cell development kinetics from PSC. PSCs were first induced into lateral plate mesoderm cells (iLPM) for 2 days. The iLPM were mixed with feeder cells (GFP^+^) to form organoid aggregates (day 2), further induced into iNK cells on transwell for 25-day induction. GFP^-^ cells from the organoids were sorted for single-cell RNA-seq on days 16, 17, 18, 19, and 20. **b,** UMAP illustrating the projection of NKP, NK with e-iNKP, l-iNKP, and im-iNK. NKP, natural NK progenitor cells, NK, natural NK cells, other-iHPC, other induced hematopoietic cells. e-iNKP, early iNK progenitor cells. l-iNKP, late iNK progenitor cells. im-iNK, immature iNK cells. **c,** pseudo-time analysis of other-iHPC, e-iNKP, l-iNKP, and im-iNK. **d,** Violin plots of gene expression levels of *CD34*, *CD38*, *CD7*, CD123-encoding *IL3RA*, CD127-encoding *IL7R*, CD10-encoding *MME*, CD56-encoding *NCAM1*, and CD16-encoding *FCGR3A* in other-iHPC, iNKP (e-iNKP and l-iNKP) and im-iNK. **e,** Flow cytometry of iNKP cells (CD45^+^CD56^-^ CD34^+^CD7^+/-^) from day-19 organoids. Representative data from three independent experiments are shown. **f,** The strategy of functional evaluation of the iNKP cells. **g,** Flow cytometry of iNK cells generated from iNKP cells *in vitro* after 14-day culture. Representative data from three independent experiments are shown. **h,** Flow cytometry of iNK cells in PB from the B-NDG hIL15 recipients 14 days after iNKP cell infusion. Representative data from three independent experiments are shown. **i,** Boxplots showing the expression levels of endogenous *CXCR4* in NKP and BM_NK cells at the transcriptome level. STRT-seq data from GSE149938 were analysed. **j,** Boxplots showing the expression levels of endogenous *CXCR4* in UCB_NK, PB_NK, iNKP, and iNK populations. Droplet-based scRNA-seq data from HRA007978 (UCB_NK) and HRA001609 (PB_NK, iNK) were analysed. **k,** Flow cytometry of the frequencies of CXCR4 in PB-NK, UCB-NK, iNKP, and iNK. Representative data from three independent experiments are shown. **l,** Amino acid sequence alignment of human CXCL12 and mouse Cxcl12. **m,** Schematic diagram of generation of CXCR4-expressing PSC (R4-PSC) and induction of CXCR4-expressing iNKP and iNK cells. **n,** Flow cytometry of the frequencies of exogenous CXCR4 in R4-PSC. **o,** Flow cytometry of the R4-iNKP cells from day-19 organoids. Representative data from three independent experiments are shown. **p,** Flow cytometry of iNK cells generated from R4-iNKP cells *in vitro* after 14 days of culture. Representative data from three independent experiments are shown. **q,** Flow cytometry analysis of iNK cells in PB from the B-NDG hIL15 recipients 14 days after R4-iNKP cell infusion. **r,s** Statistical analysis of the percentages of iNK cells in PB from the R4-iNKP cell recipients 14-, 21-, and 28-day (**r**) and 38-, 45-, 52-, and 80-day (**s**) post-infusion (n = 3 animals). **t,** Statistical analysis of the expression of CXCR4 in iNK cells. Each symbol represents an individual recipient. Data is represented as the mean ± s.d..

To facilitate the enrichment of iNKP cells, including e-iNKP and l-iNKP, for further functional evaluation and infusion purposes, we next focused on the typical markers defining natural NKP cells, including CD34, CD38, CD7, CD45RA, CD10, CD123, and CD127^23^. At the transcriptomic level, the iNKP cell population shared standard molecular features with natural NKP cell counterparts, including expressing CD34 (67.2 %), CD38 (52.7 %), and CD7 (56.8%), and rarely expressing CD123 (25%) and CD127 (23%). However, iNKP cells did not express CD10, expressed in NKP cells. The iNKP barely expressed NCAM-1 (CD56) and FCGR3A (CD16), which are typical markers of mature NK cells (**Fig. 1d**). Flow cytometry analysis of the single cells from day-19 organoids indicated an apparent CD34^+^CD7^+/-^ phenotypic progenitor cell population (≥ 79%) in the total hematopoietic cells **(Fig. 1e and Extended Data Fig. 2a)**. We sorted the CD45^+^CD56^-^CD34^+^CD7^+/-^ cells from the day-19 organoids to functionally confirm their iNK commitment potential via 14-day NK cell maturation via *in vitro* culture and infusion assays using immune-deficient mice expressing human IL15 (B-NDG hIL15) as recipients **(Fig. 1f)**. Our results showed that CD45^+^CD56^-^ CD34^+^CD7^+/-^ iNKP cells matured into CD56^+^CD16^+/-^ iNK cells with over 99% purity in culture dish within two weeks (**Fig. 1g and Extended Data Fig. 2b**). In the infusion setting, CD45^+^CD56^-^CD34^+^CD7^+/-^ iNKP cells produced dominant iNK cells co-expressing CD56 and CD16 *in vivo* 14 days post-infusion (**Fig. 1h**). However, we noticed that infusion of 2 × 10^5^ iNKP cells resulted in low chimera rates of human NK cells in the PB of B-NDG hIL15 recipients **(Fig. 1h)**.

### Enforced expression of CXCR4 promotes iNKP cell engraftment and results in abundant iNK cells persisting in PB over 80 days

SDF-1 receptor, encoded by the *CXCR4*, is an essential chemokine receptor that plays a vital role in hematopoietic stem and progenitor cell homing^26,27^. We analysed the expression levels of endogenous *CXCR4* in NKP, NK, iNKP, and iNK cell populations at the transcriptome level. Indeed, NKP and NK cells from published data (GSE149938) expressed CXCR4. Our data indicated that NK cells from umbilical cord blood (UCB) and PB also expressed *CXCR4*; however, PSC-derived iNKP and iNK cells barely expressed *CXCR4* **(Fig. 1i,j)**. Immunostaining of CXCR4 further confirmed that neither iNKP nor iNK cells expressed CXCR4 at the protein level **(Fig. 1k)**. Human CXCR4, ligand CXCL12, and mouse Cxcl12 are highly conserved except for two amino acid variations **(Fig. 1l)**. Enforced expression of human CXCR4 in murine CD4-positive T cells promotes their homing to bone marrow in mice ^28^. Thus, we propose that enforced expression of human CXCR4 in iNKP cells might promote these cells’ homing to the bone marrow of B-NDG hIL15 mice **(Fig. 1m)**. We introduced the human CXCR4 expression element at the PSC (R4-PSC) stage and successfully achieved CXCR4 expression in PSC (R4-PSC, **Fig. 1n)**, derived iNKP (R4-iNKP, **Fig. 1o**) and iNK (R4-iNK, **Fig. 1p**) cells *in vitro* (**Fig. 1, m-p**). Then, we infused 2 × 10^5^ R4-iNKP cells into individual B-NDG hIL15 recipients and weekly measured the iNK contributions in PB (**Fig. 1q**). Contrary to iNKP cell infusion producing low levels of iNK cells *in vivo*, R4-iNKP cell infusion gave rise to abundant iNK cells, with an average over 40% of total blood nucleated cells being iNK cells on day 14, over 30% on day 21, and over 20% on day 28 after infusion (**Fig. 1r and Extended Data Fig. 3a**). Despite the sharp decreases of iNK cells 28 days after R4-iNKP cell infusion, we still observed over 4% iNK cells in recipient PB 52 days post-infusion and over 0.5% iNK cells in recipient PB 80 days after infusion analysed by flow cytometry (**Fig. 1s and Extended Data Fig. 3a**). These iNK cells expressed variable but significant levels of CXCR4 *in vivo* (**Fig. 1t and Extended Data Fig. 3a**).

Collectively, we generated and characterised human pluripotent stem cell-derived NK progenitor cells *in vitro*. A single dose infusion of CXCR4-expressing iNKP cells produces abundant iNK cells *in vivo* in hIL-15-expressing recipients, which lasted over 80 days in peripheral blood.

### CD19CAR-R4iNKP cell infusion efficiently outputs CD19CAR-R4iNK cells distributed in multiple organs

Considering the lack of persistent efficacy of traditional CAR-NK cell therapy, we propose that infusion of CAR-R4iNKP cells might achieve superior therapeutic outcomes due to their potential for prolonged production of CAR-iNK cells *in vivo*. We introduced CD19CAR expressing elements into the R4-PSC to generate the CD19CAR-R4PSC line to derive CD19CAR-R4iNKP. We then assessed the CD19CAR-R4iNK cell generating potential of CD19CAR-R4iNKP cells both *in vitro* and *in vivo* (**Fig. 2a**). After introducing CD19CAR expression elements, R4-PSC expressed CD19CAR (CD19CAR-R4PSC) at a rate over 99% (**Fig. 2b**). The derived CD19CAR-R4iNKP cells were 99% CAR positive (**Fig. 2c**). A 14-day maturation assay *in vitro* confirmed that the CD19CAR-R4iNKP cells gave rise to over 99% iNK (CD19CAR-R4iNK) cells, which expressed high levels of CXCR4 (75%) and CD19CAR (> 90%) (**Fig. 2d**). We further infused 2 × 10^5^ CD19CAR-R4iNKP cells into B-NDG hIL15 mice and observed abundant CD19CAR-R4iNK cells in PB on as early as day 14 after cell infusion (**Fig. 2e**). Consistent with the patterns observed in R4-iNKP infused animals, the CD19CAR-R4iNKP cell infusion persistently gave rise to iNK cells *in vivo*, with an average over 30% of total blood nucleated cells being CD45^+^CD56^+^CD16^+/-^ on day 14, over 40% on day 21, and over 20% on day 28 (**Fig. 2f and Extended Data Fig. 4a,b**). Twenty-eight days after infusion, the ratios of CD19CAR-R4iNK cells sharply decreased to levels lower than 4% in PB (**Fig. 2g and Extended Data Fig. 4b**). Even 80 days after infusion, we still detected iNK cells in the recipient PB (> 0.3%) (**Fig. 2g and Extended Data Fig. 4b**). As expected, these *in vivo* generated iNK cells expressed high levels of CXCR4 (> 50%) and CD19CAR (> 40%) (**Fig. 2h,i and Extended Data Fig. 4b**). Next, we investigated the tissue distributions of iNK cells 21 days after CD19CAR-R4iNKP cell infusion. In the single-cell suspensions isolated from ground tissues of bone marrow (BM), liver, lung, and spleen, we observed the existence of iNK cells with a typical phenotype of CD45^+^CD56^+^CD16^+/-^, and no existence of CD3^+^ T cells, CD19^+^ B cells, or CD33^+^ Myeloid cells. These iNK cells expressed variable but significant levels of CXCR4 and CD19CAR *in vivo* (**Fig. 2j,k**).

**Fig. 2.**
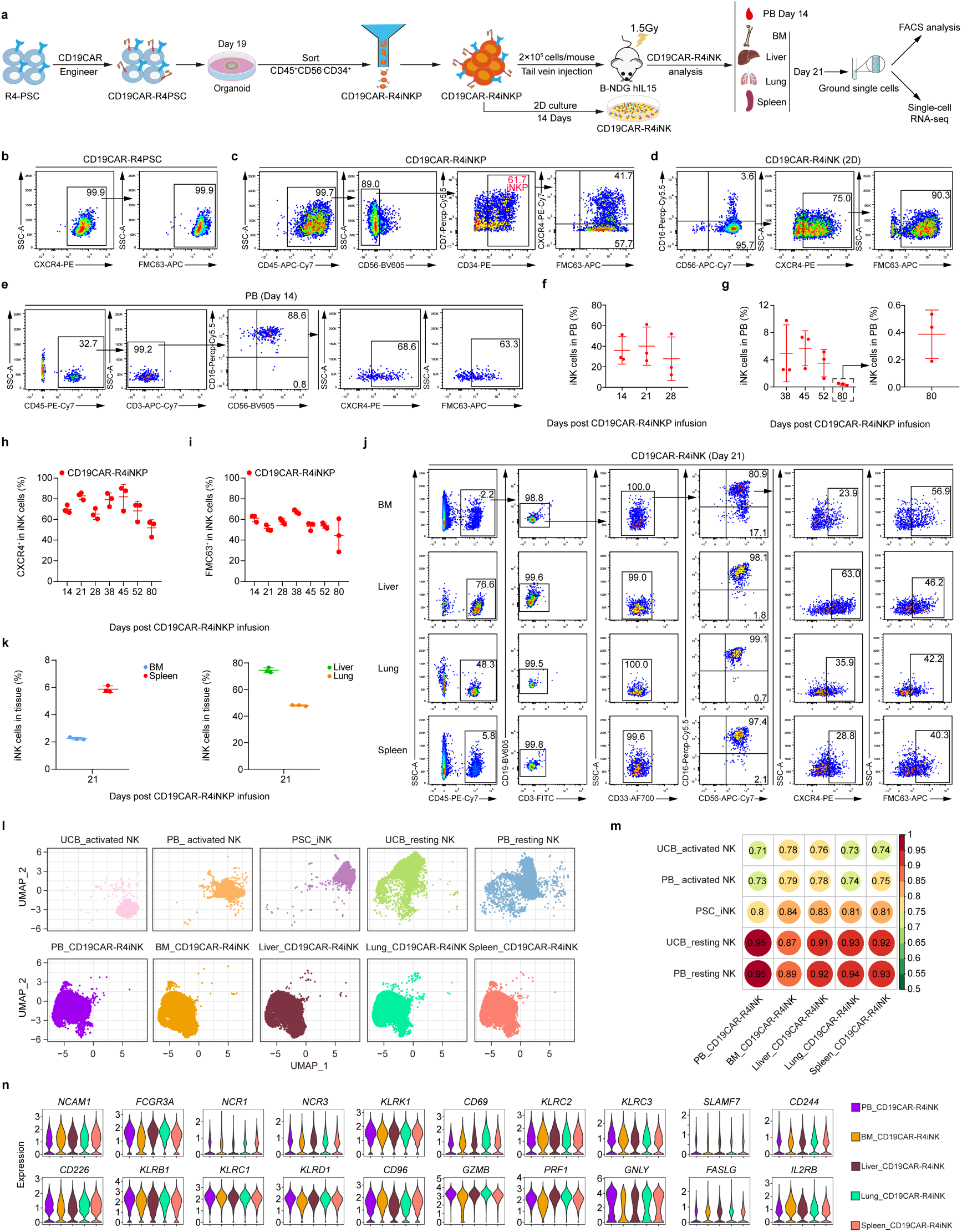
CD19CAR-R4iNKP efficiently generated homogeneous CD19CAR-iNK cells in PB and multiple organs. **a,** Strategy of functional evaluation of the CD19CAR-R4iNKP cells. **b,** Flow cytometry showing the frequencies of CXCR4 and CD19CAR (FMC63^+^) in CD19CAR-R4PSC. **c,** Flow cytometry of CD19CAR-R4iNKP cells from day-19 organoids. Representative data from three independent experiments are shown. **d,** Flow cytometry of iNK cells differentiated from CD19CAR-R4iNKP *in vitro*. Representative data from three independent experiments are shown. **e,** Flow cytometry analysis of iNK cells in PB from CD19CAR-R4iNKP cell recipients 14 days after cell infusion. **f,g**, Statistical analysis of the percentages of iNK cells in PB from CD19CAR-R4iNKP cell recipients 14-, 21-, and 28-day (**f**) and 38-, 45-, 52-, and 80-day (**g**) post-infusion (n = 3). **h,i**, Statistical analysis of the expression of CXCR4 (**h**) and CD19CAR (FMC63^+^) (**i**) in iNK cells. Each symbol represents an individual recipient. Data represents as the mean ± s.d.. **j,** Flow cytometry of iNK cells in bone marrow (BM), Liver, Lung, and Spleen of the CD19CAR-R4iNKP cell recipients (n = 3) 21 days after infusion. **k,** Statistical analysis of iNK cells in BM, Liver, Lung, and Spleen. **l,** UMAP plots showing the projections among UCB_activated NK, PB_activated NK, PSC_iNK, UCB_resting NK, PB_resting NK, and PB_CD19CAR-R4iNK, BM_CD19CAR-R4iNK, Liver_CD19CAR-R4iNK, Lung_CD19CAR-R4iNK, and Spleen_CD19CAR-R4iNK. Each tissue-derived iNK cell (CD45^+^CD56^+^) was obtained from the B-NDG hIL15 recipients 21 days after infusion of CD19CAR-R4iNKP cells, respectively. **m,** Heatmap showing the correlation value between each two populations in whole transcriptome levels. The pearson correlation coefficient presenting in each dot was encoded by color and calculated by the ‘cor’ function using the average expression of the total transcriptome profile. **n,** Violin plots of gene expression levels of CD56-encoding *NCAM1*, CD16-encoding *FCGR3A*, NKp46-encoding *NCR1*, NKp30-encoding *NCR3*, NKG2D-encoding *KLRK1*, *CD69*, NKG2C-encoding *KLRC2*, NKG2E-encoding *KLRC3*, CD319-encoding *SLAMF7*, *CD244*, *CD226*, CD161-encoding *KLRB1*, NKG2A-encoding *KLRC1*, CD94-encoding *KLRD1*, *CD96*, *GZMB*, perforin-encoding *PRF1*, *GNLY*, CD178-encoding *FASLG*, and *IL2-RB*.

Tissue-resident NK cells show phenotypic and functional heterogeneities^29^. To characterise the iNK cells produced *in vivo* in different organs, we analysed the transcriptome features of CD19CAR-R4iNKP-derived iNK cells isolated from PB, BM, liver, lung, and spleen. Contrary to the heterogeneity reported in human organ-derived NK cells^30,31^, UMAP analysis demonstrated that the iNK cells isolated from PB, BM, liver, lung, and spleen of CD19CAR-R4iNKP infused mice were highly homogeneous (**Fig. 2l**). Despite developing in xenograft animals expressing hIL-15, These *in vivo*-derived CD19CAR-R4iNK cells were closer to resting natural NK cells from UCB and PB at the global transcriptome level (**Fig. 2l,m**). In addition, the iNK cells isolated from CD19CAR-R4iNKP cell recipients’ organs expressed the typical surface marker genes of NK cells, including activating receptor gene CD56-encoding *NCAM1*, CD16-encoding *FCGR3A*, NKp46-encoding *NCR1*, NKp30-encoding *NCR3*, NKG2D-encoding *KLRK1*, *CD69*, NKG2C-encoding *KLRC2*, NKG2E-encoding *KLRC3*, CD319-encoding *SLAMF7*, *CD244*, and *CD226*, inhibitory receptors gene CD161-encoding *KLRB1*, NKG2A-encoding *KLRC1*, CD94-encoding *KLRD1*, and *CD96*, cytotoxic or survival related gene *GZMB*, perforin-encoding *PRF1*, *GNLY*, CD178-encoding *FASLG*, and *IL2-RB* (**Fig. 2n**). Collectively, infusion of iNKP cells expressing CXCR4 and CD19CAR persistently produced mature and homogenous CAR-iNK cells in multiple organs.

### CAR-R4iNKP cells mainly migrate to and develop in the bone marrow and mature into iNK cells within 14 days

To spatiotemporally trace the homing and early development of CD19CAR-R4iNKP cells in B-NDG hIL15 mice, we established the luciferase-expressing CD19CAR-R4PSC (CD19CAR-R4PSC-luci) line, and luciferase-expressing CD19CAR-PSC (CD19CAR-PSC-luci) line as homing control (**Extended Data Fig. 5a**). The derived CD19CAR-R4iNKP-luci cells and CD19CAR-iNKP-luci cells (CD45^+^CD56^-^ CD34^+^CD7^+/-^) were respectively infused into irradiated (1.5 Gy) B-NDG hIL15 mice via tail vein injection (n = 4 in each group, 1 × 10^5^ cells/mouse). We performed Bioluminescence imaging (BLI) assays at indicated time points post-infusion to trace the spatiotemporal homing sites of CD19CAR-iNKP cells and organ distributions of their subsequent progenies. Meanwhile, we analysed the kinetics of iNK cells in PB (**Fig. 3a**). BLI of the body’s dorsal, left, and ventral views demonstrated that the CD19CAR-R4iNKP-luci cells and their developing progenies were broadly distributed in the femur, spinal cord, skull, and sternum bone marrow, and cervical lymph nodes on day 5 after infusion. However, the CD19CAR-iNKP-luci-infused mice barely showed fluorescence signals on day 5, indicating that the enforced expression of CXCR4 promoted the iNKP homing to bone marrow (**Fig. 3b**). The kinetics of spinal cord luminescence signaled that iNKP developed mainly in the bone marrow and seemingly proliferated to a peak on day 11 and then sharply declined (**Fig. 3c**). On day 14, the CD19CAR-R4iNKP-luci progeny cells appeared in the recipients’ liver, lung, and spleen (**Fig. 3b**). Total luminescence kinetics indicated that iNK cells accumulated to a peak on day 21 measured from the dorsal, left, and ventral views (**Fig. 3, d-f**). The values of total flux sharply increased from day 5 to day 21, then decreased from day 21 to day 28 in the CD19CAR-R4iNKP-luci cell recipients. However, the CD19CAR-iNKP-luci cell recipients showed extremely low radiance from day 5 to day 28 (**Fig. 3b,d-f**). Notably, over 10% of iNK cells already emerged in the PB of CD19CAR-R4iNKP-luci cell recipients on day 14 (**Fig. 3g and Extended Data Fig. 5b**). The highest levels of iNK output in PB seemingly lasted for around one week, then declined starting from day 21, consistent with the signal peaks of the total flux values. However, iNK cells were barely detected in the PB of CD19CAR-iNKP-luci cell recipients (**Fig. 3g and Extended Data Fig. 5b**). These results suggest that CXCR4 promotes iNKP cell migration to bone marrow. The CAR-R4iNKP cells mainly develop in the bone marrow and mature into iNK cells within 14 days.

**Fig. 3.**
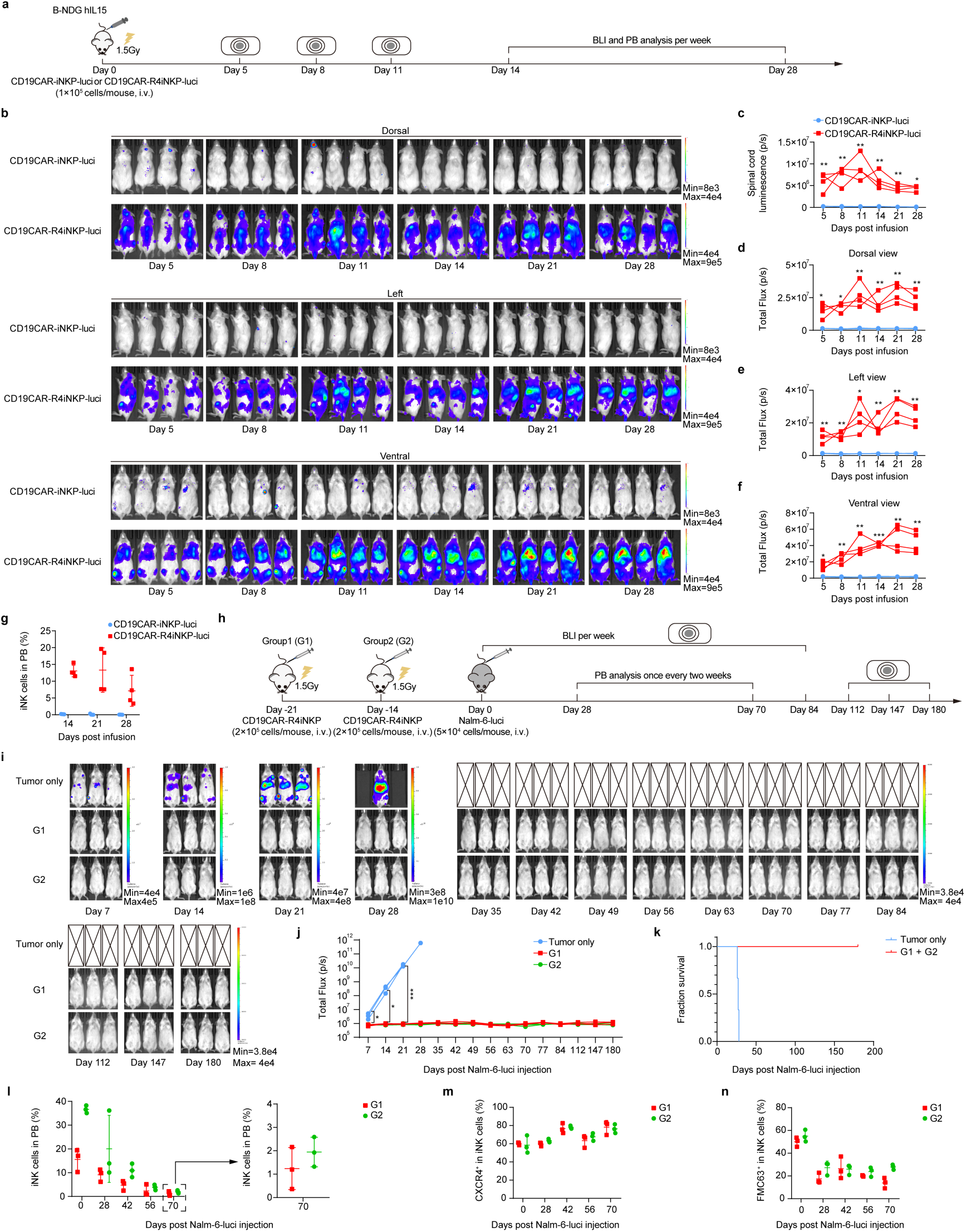
CD19CAR-R4iNKP cells migrate to and develop in bone marrow, preventing animal tumours. **a,** Schematic diagram of evaluating the distributions of CD19CAR-R4iNKP and their progeny cells in B-NDG hIL15 recipients after infusion. **b,** BLI images (Dorsal, Left, and Ventral) showing the distributions of iNKP and their progenies of mice infused with CD19CAR-iNKP-luci or CD19CAR-R4iNKP-luci. **c,** Statistical analysis of the spinal cord luminescence (p/s) of the CD19CAR-iNKP-luci cell recipients or CD19CAR-R4iNKP-luci cell recipients. **d,** Statistical analysis of the total flux (p/s) of the Dorsal view of the CD19CAR-iNKP-luci cell recipients or CD19CAR-R4iNKP-luci cell recipients. **e,** Statistical analysis of the total flux (p/s) of the Left view of the CD19CAR-iNKP-luci cell recipients or CD19CAR-R4iNKP-luci cell recipients. **f,** Statistical analysis of the total flux (p/s) of Ventral view of the CD19CAR-iNKP-luci cell recipients or CD19CAR-R4iNKP-luci cell recipients. **g,** Statistical analysis of iNK cells in PB of the CD19CAR-iNKP-luci cell recipients or CD19CAR-R4iNKP-luci cell recipients on days 14, 21, and 28 post-infusions. **h,** Schematic diagram of evaluating tumour prevention potential of the CD19CAR-R4iNKP cell recipients. Tumour only group: mice were only injected with Nalm-6-luci cells, G1 (Group 1): mice were infused with CD19CAR-R4iNKP cells on day −21 and injected with Nalm-6-luci cells on day 0. G2 (Group 2): mice were infused with CD19CAR-R4iNKP cells on day −14 and injected with Nalm-6-luci cells on day 0. **i,** BLI of the human-tumour-injected mice from tumour only group, G1 and G2 (n = 3 mice per group) on days 7, 14, 21, 28, 35, 42, 49, 56, 63, 70, 77, 84, 112, 147, and 180. **j,** Statistical analysis of the tumour-injected mice’s total flux (p/s). **k,** Kaplan-Meier survival curves of the tumour only and G1 + G2 groups. n = 3 mice for tumour only group. n = 6 mice for G1 and G2 combined group. **l,** Statistical analysis of the percentages of iNK cells in PB from CD19CAR-R4iNKP cell recipients 0-, 28-, 42-, and 56-days and 70-day post-Nalm-6-luci injection (n = 3). **m,n**, Statistical analysis of the expression of CXCR4 (**m**) and CD19CAR (FMC63^+^) (**n**) in iNK cells. Each symbol represents an individual recipient. Data is represented as the mean ± s.d.. All data represent mean ± s.d. and are analysed by two-sided unpaired Student’s t-test (**c-f**) and one-way ANOVA test (**j**). * *p* < 0.05, *** *p* < 0.001.

### CD19CAR-R4iNKP cell infusion achieves precise prevention of human B-ALL tumours in xenograft animals

Since iNKP cell infusion results in prolonged iNK cell persistence *in vivo* over 80 days, we investigated whether iNKP cell therapy could achieve precise tumour prevention. To establish CAR-iNK reconstituted immune surveillance, we infused 2 × 10^5^ CD19CAR-R4iNKP cells into B-NDG hIL15 mice. To simulate natural tumour occurrence, we respectively injected 5 × 10^4^ CD19^+^ tumour cells (Nalm-6-luci) into the animals at 2 weeks (Group 2, G2) and 3 weeks (Group 1, G1) after CD19CAR-R4iNKP cell infusion. A group of mice without the CD19CAR-R4iNKP cell infusion was injected with the same amount of tumour cells as tumour only group. BLI was performed weekly to trace the kinetics of tumour growth. For tumour burden analysis, we defined the time of tumour injection as Day 0 **(Fig. 3h)**. Notably, live imaging results showed no significant tumour signals detected in the G1 and G2 mice over 180 days. In contrast, the tumour only group developed severe malignancy and died within 28 days (Median = 27 days) **(Fig. 3i-k)**. To confirm the presence of CD19CAR-R4iNK cells within the G1 and G2 mice, we measured their PB on day 0, day 28, day 42, day 56, and day 70 by flow cytometry. The CD19CAR-R4iNK cell chimera rates on day 0 were 15.6% ± 4.8% in G1 mice and 36.6% ± 1.6% in G2 mice **(Fig. 3l and Extended Data Fig. 6a,b)**. The CD19CAR-R4iNK cells gradually decreased to 1.2% ±0.9% in G1 mice and 1.9% ± 0.6% in G2 mice on day 70 **(Fig. 3l, and Extended Data Fig. 6a,b)**. The expression levels of exogenous CXCR4 in CD19CAR-R4iNK cells were stable **(Fig. 3m and Extended Data Fig. 6a,b)**. Additionally, we examined the dynamic changes of CD19CAR expression in CD19CAR-R4iNK cells. The CD19CAR expression positive rates were 50.0% ± 4.1% in G1 and 55.0% ± 5.2% in G2 on day 0 and then gradually declined to 13.9% ± 4.6% in G1 and 27.9% ± 2.3% in G2 on day 70 **(Fig. 3n and Extended Data Fig. 6a,b)**. These results indicate that CAR-iNKP cell therapy can achieve precise tumour prevention.

### Chemotherapy combined with CD19CAR-R4iNKP cell therapy achieves long-term remission in animals bearing human B-ALL tumours

Chemotherapy is a conventional approach broadly used to treat human tumour patients. The residual tumour cells after chemotherapy are the major causes of tumour relapse ^32,33^. We investigated whether combining chemotherapy and CD19CAR-R4iNKP cell infusion could achieve prolonged complete remission in animals bearing human B-ALL tumour cells. We intravenously injected the Nalm-6-luci cells (1 × 10^5^ cells/mouse) into the B-NDG hIL15 mice to generate a human B-ALL xenograft model. The tumour cell injected mice were randomly divided into three groups: untreated (tumour only, n = 4), treated with cyclophosphamide (Chemo, n = 4), and treated with cyclophosphamide and CD19CAR-R4iNKP (Chemo + CD19CAR-R4iNKP, n = 4). We started a chemotherapy regimen with cyclophosphamide (150 mg/kg). Three days later, the Chemo + CD19CAR-R4iNKP group mice were intravenously injected with CD19CAR-R4iNKP cells (5 × 10^5^ cells/mouse). We performed BLI every seven days to monitor the mice’s tumour burden. The kinetics of CD19CAR-R4iNK cells in PB of the Chemo + CD19CAR-R4iNKP group were analysed once a week from day 17-45 by flow cytometry (**Fig. 4a**). In the early stage (Day 7-14), the Chemo group and Chemo + CD19CAR-R4iNKP group exhibited similar tumour burden trends, mainly attributing to the therapeutic outcome of Chemo. However, the tumour burden of the Chemo group gradually aggravated starting from day 14 (**Fig. 4b,c**). We had to sacrifice the mice due to paralysis caused by heavy tumour burden from the tumour only group during days 19-21 (Median = 19.5) and from the Chemo group during days 36-51 (Median = 46) (**Fig. 4b-d**). Notably, the mice showed complete remission over 150 days after Chemo and CD19CAR-R4iNKP combined treatment (**Fig. 4b-d and Extended Data Fig. 7a**). Consistent with the long-term remission, the CD19CAR-R4iNK cells in PB of Chemo + CD19CAR-R4iNKP group animals were still detected (4.2% ± 1.9%) by flow cytometry on day 45 (**Fig. 4e,f and Extended Data Fig. 8a**). The CD19CAR-R4iNK cells sustainably expressed high levels of CXCR4 and CD19CAR on their surfaces (**Fig. 4e,g,h and Extended Data Fig. 8a**). These results show that CD19CAR-R4iNKP cell infusion has eliminated the MRD and achieved long-term complete remission.

**Fig. 4.**
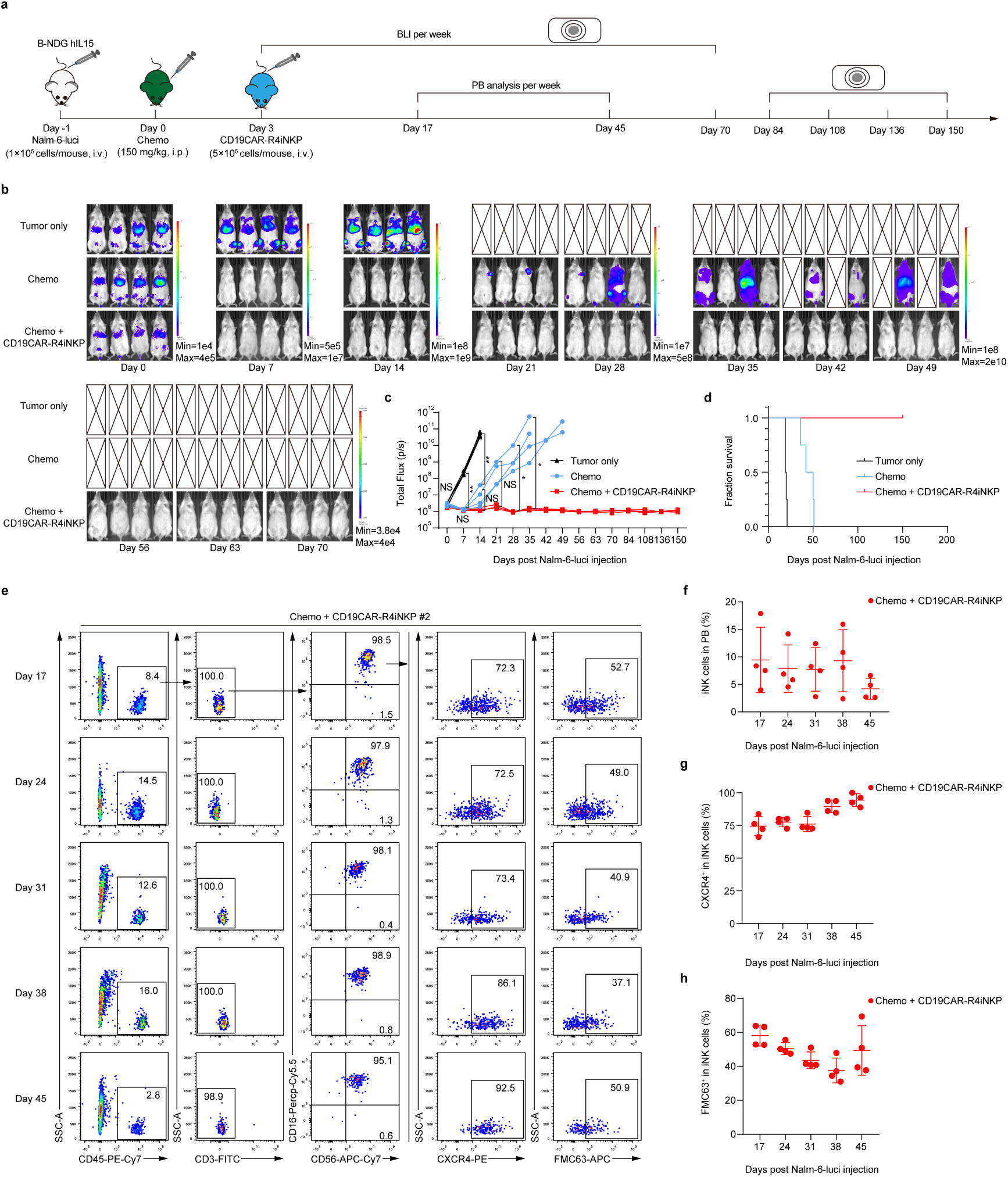
CD19CAR-R4iNKP cell therapy eliminates the tumour MRD, achieving long-term complete remission. **a,** Schematic diagram of evaluating the anti-tumour potential of the CD19CAR-R4iNKP in mice bearing tumour MRD. **b,** BLI of the tumour-bearing mice from tumour only group, Chemo group, and Chemo + CD19CAR-R4iNKP group (n = 4 mice per group) on days 0, 7, 14, 21, 28, 35, 42, 49, 56, 63, and 70. **c,** Kinetics of the tumour-bearing mice’s total flux (p/s) from tumour only, Chemo treated, and Chemo plus CD19CAR-R4iNKP treated animals (n = 4 animals in each group). **d,** Survival curves of the three groups of tumour-bearing mice. n = 4 mice in each group. **e,** Flow cytometry of iNK cells in PB from the Chemo + CD19CAR-R4iNKP treated animals. PB samples were collected on days 17, 24, 31, 38, and 45. **f,** Statistical analysis of iNK cells in PB of the Chemo + CD19CAR-R4iNKP treated animals on days 17, 24, 31, 38, and 45. **g,h**, Statistical analysis of CXCR4 (**g**) and CD19CAR (FMC63^+^) (**h**) in iNK cells. Each symbol represents an individual mouse. All data represent mean ± s.d. and are analysed by one-way ANOVA test (**c**), two-sided unpaired Student’s t-test (**c**), and the log-rank test (**d**). NS, not significant, * p < 0.05, ** p < 0.01.

## Discussion

NK and CAR-NK cell therapy lack sustained efficacy due to the short-term persistence of mature NK cells after infusion. In this study, we overcome the persistence bottleneck by infusing iNKP cells derived from pluripotent stem cells. Enforced expression of CXCR4 significantly promotes the engraftment of iNKP cells and produces abundant iNK cells in PB lasting over 80 days. Notably, CAR-R4iNKP cell infusion successfully gives rise to CAR-iNK cells, which precisely prevent human tumour occurrence. Chemotherapy combined with CAR-iNKP cell therapy eradicates MRD and leads to long-term complete remission in human tumour-bearing animals. For the first time, we developed *de novo* NKP cell therapy, which potentially prevents human tumour occurrence and clears MRD in combination with conventional chemotherapy.

Natural NKP cells and NK cells express high levels of CXCR4. However, endogenous CXCR4 expression in PSC-derived iNKP cells and iNK cells in this study is significantly rare, consistent with those produced via the EB and organoid methods, with only 2-11% iNK cells expressing endogenous CXCR4 ^13,34^. iNKP engraftment efficiency is extremely low without enforced exogenous CXCR4 expression, indicating that CXCR4 is essential for iNKP homing to the appropriate developmental microenvironment in the body. Consistent with our observation, the absence of Cxcr4 impairs NK cell maturation *in vivo*^35^. Future efforts should explore induction conditions to activate endogenous CXCR4 expression to avoid relying on exogenous CXCR4 for iNKP engraftment. Notably, the high conservation of CXCL12 chemokine between mice and humans ensures that mouse Cxcl12 can interact effectively with human CXCR4, allowing CXCR4-expressing iNKP cells and CAR-iNKP cells to home to developmental sites *in vivo*.

CAR-HSPCs, when introduced *in vivo*, can give rise to CAR-NK, CAR-macrophages (CAR-Mac), and CAR-B cells^36^. CAR expression in these diverse cell types could result in altered tissue distribution of multiple immune cells, raising potential unknown risks. In contrast, CAR-iNKP cell infusion into mice prominently produced CAR-iNK cells, with no evidence of other lineages, indicating that CAR-iNKP differentiation towards the NK lineage is committed. The introduction of CAR at the stem cell stage, targeting antigens like CD19 or Mesothelin, does not impact NK cell maturation *in vitro*^37–39^. This feature is advantageous for introducing various CAR constructs at the stem cell level to generate CARs-iNKP and testing their therapeutic potential against broad cancers. No CD3^+^ T cells were detected within three months in iNKP cell-infused recipients. Since the mouse thymus is a xenogeneic organ for human cells, it could obscure the potential for iNKP cells to migrate to the thymus and develop into iT cells. Human CD34^+^ HSPCs, however, have been shown to produce a small number of human T cells approximately eight weeks post-infusion ^40^. Therefore, it cannot be ruled out that iNKP cells might give rise to human T cells following infusion into patients. We confirmed that CAR-iNKP cells failed to produce CAR-iT cells when cultured *in vitro* using the conventional organoid method (unpublished data). This suggests that CAR signaling might inhibit or disrupt T cell development and maturation. Hence, clinical infusion of CAR-iNKP cells can cleverly avoid the potential risks of generating CAR-iT cells. Given that some CAR-iNKP cells might silence CAR expression during *in vivo* differentiation, it could be prudent to employ RAG1/2 gene knockout at the stem cell stage to block T cell development pathways, thereby eliminating the risk of generating any iT cells.

CD16 is an important molecule mediating ADCC and acquired immune function in NK cells. Nearly 100% of natural NK cells derived from human tissues express CD16. However, iNK cells generated by conventional induction methods, including monolayers and EB methods, exhibit only 1-20% CD16 expression^41,42^. Our previous research demonstrated that an organoid induction system increased the CD16 expression levels in iNK cells to 40-80%^13^. Unlike the *in vitro* induction system, iNKP cells derived from the same PSC background expressed nearly 100% CD16 *in vivo*, similar to natural tissue-derived NK cells. This suggests that the natural physiological microenvironment is critical for regulating CD16 expression. Thus, CD16 likely serves as a key indicator of normal NK cell maturation.

In this study, we observed that iNKP cell infusion produced iNK cells distributed in multiple organs, including bone marrow, liver, lung, and spleen. Whether iNKP cells can independently develop in multiple mouse organs remains uncertain. Natural human NK cells follow independent developmental pathways in different organs, showing functional heterogeneity^43–46^. However, iNK cells derived from iNKP cell infusion in mice lacked this heterogeneity. This could be due to differences in developmental potential between *in vitro*-induced iNKP cells and natural NKP cells, specifically the inability of iNKP cells to generate more diverse NK subtypes. Alternatively, the xenogeneic environment of mouse organs may not provide the conditions for the independent development and functional specialization seen in human organ-derived NK cells. Therefore, iNKP cell infusion into humans might exhibit organ-specific NK development and heterogeneity. iNK cells generated *in vivo* from iNKP cells sustained for 80 days and then largely vanished. Since NKP cells lack self-renewal capabilities, they can only proliferate and differentiate into NK cells post-infusion. This characteristic ensures that iNKP and CAR-iNKP cell infusion are safe, with significantly reduced risks of long-term production and potential side effects associated with iNK or CAR-iNK cells.

Upon CAR-iNKP cell infusion, mature CAR-iNK cells were produced within 14 days *in vivo* and successfully resisted artificially introduced tumours, demonstrating their effective preventive capabilities. This finding has profound implications for the early prevention of cancers. By designing multiple tumour targets into CAR constructs, infusing CARs-iNKP cells could prevent various tumour occurrences. In this study, we only conceptually tested the effectiveness of CAR-iNKP cell infusion for preventing CD19^+^ tumours, and further testing with multiple tumour targets is needed to assess its broader tumour prevention potential.

Combined chemotherapy and CAR-iNKP cell infusion achieved long-term complete remission for up to five months in mice with CD19^+^ B-cell tumours. Further observation is required to determine whether a complete cure has been achieved. This approach suggests that CAR-iNKP cells can rapidly generate CAR-iNK cells after infusion during periods of low tumour burden following chemotherapy, effectively clearing residual tumours. This implies that in clinical scenarios involving tumour MRD-positive cases after chemotherapy, combining CAR-iNKP cell therapy might achieve MRD-negative transformation.

Further testing of the clinical translational potential of iNKP and CAR-iNKP will need to address compatibility issues with patient immunity. Autologous iPSC-derived CAR-iNKP for tumour prevention and treatment could minimize immune rejection due to reduced immunogenicity. In the case of allogeneic CAR-iNKP, strategies to improve immune compatibility are required, such as gene editing at the stem cell stage to reduce host immune rejection of allogeneic CAR-iNKP and CAR-iNK cells. Pre-treatment regimens similar to clinical allogeneic immune cell therapies might also enhance the persistence of CAR-iNKP cells post-infusion.

In conclusion, we identify a *de novo* iNKP cell population induced from pluripotent stem cells, which can produce abundant and persistent iNK cells *in vivo* after infusion. CAR-iNKP cell infusion alone effectively prevents tumour occurrence. Combined with chemotherapy, CAR-iNKP cell infusion led to long-term remission in human B-ALL tumour-bearing animals. This study offers a valuable reference for developing iNKP and CAR-iNKP cell therapy for tumour prevention and eradication of MRD in cancer patients.

## Methods

### Mice

B-NDG hIL15 (NOD.CB17-Prkdc^scid^ Il2rg^tm1Bcgen^ Il15^tm1(IL15)^ ^Bcgen^/Bcgen) mice were purchased from Biocytogen. Mice were housed in the SPF-grade animal facility of the Institute of Zoology, Chinese Academy of Sciences.

### Cell culture

The PSC line (Q380-ESC) was provided by the National Stem Cell Resource Center, Institute of Zoology, Chinese Academy of Sciences. The PSC differentiation toward iNK differentiation and the related anti-tumour activity assessments of iNK cells in animals are approved by the Biomedical Research Ethics Committee of the Institute of Zoology, Chinese Academy of Sciences. The Q380-ESC and derived R4-PSC line, CD19CAR-R4PSC line, CD19CAR-PSC-luci line, and CD19CAR-R4PSC-luci line were maintained in Essential 8 medium (Gibco, cat.no. A1517001) on vitronectin (Gibco, cat.no. A31804) coated plates. The OP9 cell line was purchased from ATCC and cultured with α-MEM (Gibco, cat.no. 1251063) with 20% fetal bovine serum (FBS) (Ausbian, cat.no. WS50T). The luciferase-expressing Nalm-6 cells, kindly provided by Professor Min Wang (Leukemia Center, Institute of Hematology and Blood Diseases Hospital, Chinese Academy of Medical Sciences, Tianjin, China), were cultured in RPMI 1640 medium (Gibco, cat. no. C11875500BT) supplemented with 10% FBS. UCB samples were obtained from the Guangdong Cord Blood Bank (Guangzhou, China).

### Cell line engineering

The cDNA encoding *CXCR4* was inserted into the PiggyBac expression vector (PB530A-2, SBI) to generate a recombinant vector. The *CXCR4*-expressing PiggyBac vector and the transposase expression vector were electroporated into PSC using Electroporator EX+ (Celetrix, 11-0106). Seven days after electroporation, PSCs were sorted for two rounds to establish the R4-PSC. For the CD19CAR-R4PSC construction, the CD19 CAR construct (FMC63 scFV-CD8α hinge-CD8α TM-CD3ζ) was inserted into the PiggyBac expression vector to generate the PB-CD19 CAR vector. Then, the PB-CD19 CAR vector was electroporated into R4-PSC with the transposase expression vector. CD19CAR-positive PSC was sorted for two rounds to establish the CD19CAR-R4PSC after seven days of electroporation. To construct the luciferase-expressing CD19CAR-PSC and CD19CAR-R4PSC, luciferase-P2A-puro elements (luciferase encoding sequences-P2A-puromycin-resistance (puro) sequences) were inserted into PiggyBac expression vector (PB530A-2, SBI) which was electroporated into CD19CAR-PSC and CD19CAR-R4PSC. The puro-resistance clones were picked to establish the PSC lines of CD19CAR-PSC-luci or CD19CAR-R4PSC-luci.

### iNKP induction

The method for hematopoietic differentiation of PSC has been previously described^13^. Firstly, PSC, R4-PSC, CD19CAR-R4PSC, CD19CAR-PSC-luci, and CD19CAR-R4PSC-luci were respectively plated in 10 cm Petri dishes(Greiner, 664160) for lateral plate mesoderm (iLPM) cell induction. After a 2-day induction, iLPM cells were harvested. Then, 2 × 10^4^ iLPM cells were combined with 5 × 10^5^ GFP^+^OP9 cells to form organoid aggregates and seeded onto a transwell insert (LABSELECT,14111) for iNKP generation. After 17-day induction on the air-liquid interface, CD45^+^CD56^-^ CD34^+^CD7^+/-^ iNKP were sorted by FACS.

### Generation of iNK cells *in vitro*

For iNK cell differentiation *in vitro* from iNKP by the monolayer method, Day 19 CD45^+^CD56^-^CD34^+^CD7^+/-^ iNKP were sorted by FACS and cultured in a T25 culture flask(NEST, 707003)at a density of 1 × 10^6^ cells/ml using NK differentiation medium^47^. Complete medium exchanges were performed every two days. Mature iNK cells were harvested after 14 days.

### iNK and CAR-iNK generation *in vivo*

B-NDG hIL15 mice were irradiated (1.5 Gy) by X-ray (Rad Source, RS2000). To evaluate the role of CXCR4 in promoting homing and the capability of producing R4-iNK and CD19CAR-R4iNK *in vivo* from R4-iNKP and CD19CAR-R4iNKP, the iNKP, R4-iNKP, and CD19CAR-R4iNKP (2 × 10^5^ cells/mouse) were injected into the tail vein of the mice 4 hours post-irradiation. For tracing CD19CAR-R4iNKP early homing and kinetic analysis of iNK at total body level, CD19CAR-iNKP-luci and CD19CAR-R4iNKP-luci (1 × 10^5^ cells/mouse) were injected into the tail vein of the mice 4 hours post-irradiation. Mice were fed with trimethoprim-sulfamethoxazole-treated water for two weeks to prevent infection.

### Flow cytometry and Antibodies

R4-PSC, CD19CAR-PSC and CD19CAR-R4PSC, iNKP, R4-iNKP and CD19CAR-R4iNKP from day-19 organoids, iNK, R4-iNK and CD19CAR-R4iNK from 14-day culture *in vitro*, iNK, R4-iNK and CD19CAR-R4iNK from infused mice were stained with the indicated antibodies. Antibodies used in this study: Human TruStain FcX^TM^ (Biolegend, clone 422302), CD45 (Biolegend, clone HI30), CD3 (Biolegend, clone UCHT1), CD16 (Biolegend, clone 3G8), CD19 (Biolegend, clone HIB19), CD33 (Biolegend, clone WM53), CD56 (Biolegend, clone HCD56), CD7 (Biolegend, clone CD7-6B7), CD34 (Biolegend, clone 581), CXCR4 (Biolegend, clone 12G5) and anti-FMC63 scFv (BioSwan, clone M19H). The stained cells were analysed by BD LSRFortessa™ or sorted by BD FACSAria™ Fusion (BD Biosciences). R4-PSC were defined as CXCR4^+^, CD19CAR-PSC were defined as FMC63^+^, CD19CAR-R4PSC were defined as CXCR4^+^FMC63^+^, iNKP were defined as CD45^+^CD56^+^CD34^+^CD7^+/-^, R4-iNKP were defined as CD45^+^CD56^+^CD34^+^CD7^+/-^CXCR4^+^, CD19CAR-R4iNKP were defined as CD45^+^CD56^+^CD34^+^CD7^+/-^CXCR4^+^FMC63^+^, iNK were defined as CD45^+^CD56^+^CD16^+/-^, R4-iNK were defined as CD45^+^CD56^+^CD16^+/-^CXCR4^+^, CD19CAR-R4iNK were defined as CD45^+^CD56^+^CD16^+/-^CXCR4^+^FMC63^+^. The flow cytometry data was analysed by FlowJo software (version 10.8.1, Tree Star, Ashland, OR, USA).

### CAR-iNKP therapy for tumour prevention

To reconstitute CAR-iNK immune surveillance, the CD19CAR-R4iNKP cells (2 × 10^5^ cells/mouse) were sorted and infused into irradiated (1.5 Gy) B-NDG hIL15 mice (8-10 weeks, female). To challenge the CAR-iNK immune surveillance, Nalm-6-luci cells (5 × 10^4^ cells/mouse) were intravenously injected into the mice 14 (G1) or 21 (G2) days post CD19CAR-R4iNKP cell infusion, Bioluminescent Imaging (BLI) was performed at indicated times.

### Combination of chemotherapy and CAR-iNKP cell infusion

To establish the B-cell leukemia xenograft models, B-NDG hIL15 mice were intravenously injected with Nalm-6-luci (1 × 10^5^ cells/mouse) on day −1. On day 0, the tumour-bearing mice were randomly divided into three groups (tumour only, Chemo, Chemo + CD19CAR-R4iNKP, n = 4 mice in each group). Mice in the Chemo group and Chemo + CD19CAR-R4iNKP group were treated with Chemo (Cyclophosphamide, 150 mg/kg, i.p.) on day 0; Mice in the Chemo + CD19CAR-R4iNKP group were infused with CD19CAR-R4iNKP (5 × 10^5^ cells/mouse) on day 3. Bioluminescent Imaging (BLI) was performed at the indicated times.

### Single-cell RNA-sequencing and data analysis

For droplet-based single-cell RNA-sequencing (10× genomics), resting NK (CD45^+^CD3^-^CD56^+^) from fresh umbilical cord blood, PSC-derived cells (GFP^-^) from the day 16-20 organoids, and CD19CAR-R4iNK cells (CD45^+^CD3^-^CD56^+^) from peripheral blood, bone marrow, liver, lung and spleen of the CD19CAR-R4iNKP cell recipients were sorted respectively. The scRNA-seq data were generated using a Chromium system (10× Genomics, PN120263) following the manufacturer’s instructions. Each scRNA-seq datasets were subsequently aligned and quantified utilizing the CellRanger software package (version 7.0.1) with GRCh38 human reference genome (https://cf.10xgenomics.com/supp/cell-exp/refdata-gex-GRCh38-2020-A.tar.gz). Then, datasets were aggregated by the ‘aggr’ function of the CellRanger package and subjected to Seurat (version 4.2.0)^48^ for further analysis.

PSC-derived cells (Day 16-20 organoids) with gene numbers ranging from 1,000 to 10,000 and reads mitochondrial gene aligned below 10% were retained (54,624 cells) for principal component analysis (PCA), cell clustering and uniform manifold approximation and projection (UMAP) dimensional reduction analysis. The top 2,000 highly variable genes were used to perform PCA. The 30 most variable PCs were selected to accomplish the UMAP analysis and cell clustering with a resolution = 0.3. Six cell populations were identified and annotated by typical cell markers. For determining the iNKP cells derived from PSC, we extracted the cells co-expressing PTPRC (CD45) and CD34 and the iNK lineage population for re-clustering (nPC = 30, resolution = 0.6). Four cell populations were identified and annotated by integration analysis with natural NKP and NK (GSE149938). To illustrate the development trajectory of iNKP and im-iNK cells, we performed pseudo-time analysis with the Monocle 3 package^49^. The violin plots exhibited the expression distribution implemented by ‘Vlnplot’ of Seurat and ggplot2 (v 4.2.0).

The scRNA-seq data of the single cells of CD19CAR-R4iNKP cell recipients’ PB and organs were filtered with the following QC: less than 8,000 genes detected, more than 500 read counts, and less than 40,000 read counts, and less than 10% of aligned reads mapping to mitochondrial genes. The scRNA-seq data of the single cells of UCB_activated NK, PB_ activated NK, PSC-iNK, UCB_resting NK, and PB_ resting NK were filtered with the following QC: less than 8,000 genes detected, and less than 15% of aligned reads mapping to mitochondrial genes. The data were merged and applied for PCA and UMAP (nPCs = 40) analysis. The correlation between each two populations in whole transcriptome levels was calculated by the “cor” function and assessed by the pearson coefficient using the average expression of the total transcriptome profile. The heatmap was drawn by corrplot and ggplot2 package (v 3.4.2).

### Statistics

All quantitative analyses were performed with SPSS (version 21, IBM Corp., Armonk, NY, USA). The Shapiro–Wilk normality test assessed data distribution. The paired-sample *t*-test was used to compare two groups of data. The one-way ANOVA test was applied to compare three groups of data. A log-rank (Mantel-Cox) test was used for the Kaplan-Meier survival curves analysis. Statistical analysis were performed using GraphPad Prism (8.0.1, GraphPad Software).

## Ethics statement

Experiments and handling of mice were conducted under the Institutional Animal Care and Use Committee of the Institute of Zoology, Chinese Academy of Sciences. PB samples were obtained from a healthy donor with the donor’s informed consent.

## Data availability

The scRNA-seq data of UCB_resting NK cells, PSC-derived cells from day 16-20 organoids, and CD19CAR-R4iNK cells from PB, BM, Liver, Lung, and Spleen have been deposited in the Genome Sequence Archive public database (HRA008662). The scRNA-seq data of natural NKP and NK are from the Gene Expression Omnibus database (GSE149938). The accession number scRNA-seq data of UCB_activated NK is HRA007978. The accession number scRNA-seq data of PSC-iNK, PB_resting NK cells, and PB_activated cells is HRA001609.

## Code availability

R scripts with .Rmd files used to analyse scRNA-seq data have been deposited at Github and are publicly available as of the publication date. GitHub repository: https://github.com/lyqbiocc/scRNA-seq-of-PSC-iNKP.

## Acknowledgments

This work was supported by grants from the National Natural Science Foundation of China (82450001, 81925002, 82300132, 32300676) and the National Key R&D Program of China (2024YFA1108302). This research project also received support from the Hainan Lecheng Real-World Research Institute.

## Author contributions

J.Y.W. conceived and supervised the study. F.X.H. and J.Y.W. designed the experiments. Z.Q.W., F.X.H., D.H.H., L.Q.Z., C.X.X., Y.H.L., T.J.W., M.Y.Z., J.X.W. Y.Q.L., H.M.Q., L.J.L., Y.Y.S., Y.C. and Y.P.Z. performed the experiments. Z.Q.W. and F.X.H. analysed and interpreted data. Y.Q.L. and Q.T.W. analysed the RNA-seq data. J.Y.W., F.X.H., Z.Q.W., Y.Q.L., D.H.H., C.X.X., T.J.W., and M.Y.Z. wrote the manuscript. J.Y.W. provided the final approval of the manuscript.

## Competing interests

The authors declare no competing interests.

---

**Extended Data Fig. 1.**
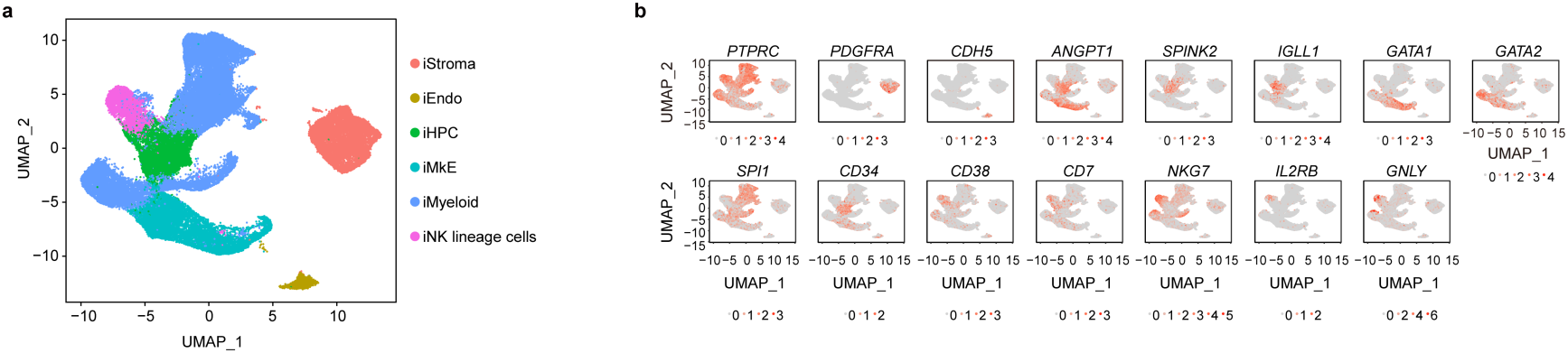
Definition of cell subpopulations based on typical markers. **a,** UMAP clustering of all the GFP^-^ cells of the organoids from days 16, 17, 18, 19, and 20. iStroma, induced Stroma cells. iEndo, induced Endothelial cells. iHPC, induced Hematopoietic progenitor cells. iMkE, induced Megakaryocytic–Erythroid cells, iMyeloid, induced Myeloid cells, iNK lineage cells, induced iNK progenitor cells and iNK cells. **b,** UMAP visualization of the feature genes expressed in iStroma, iEndo, iHPC, iMkE, iMyeloid, and iNK lineage cells.

**Extended Data Fig. 2.**
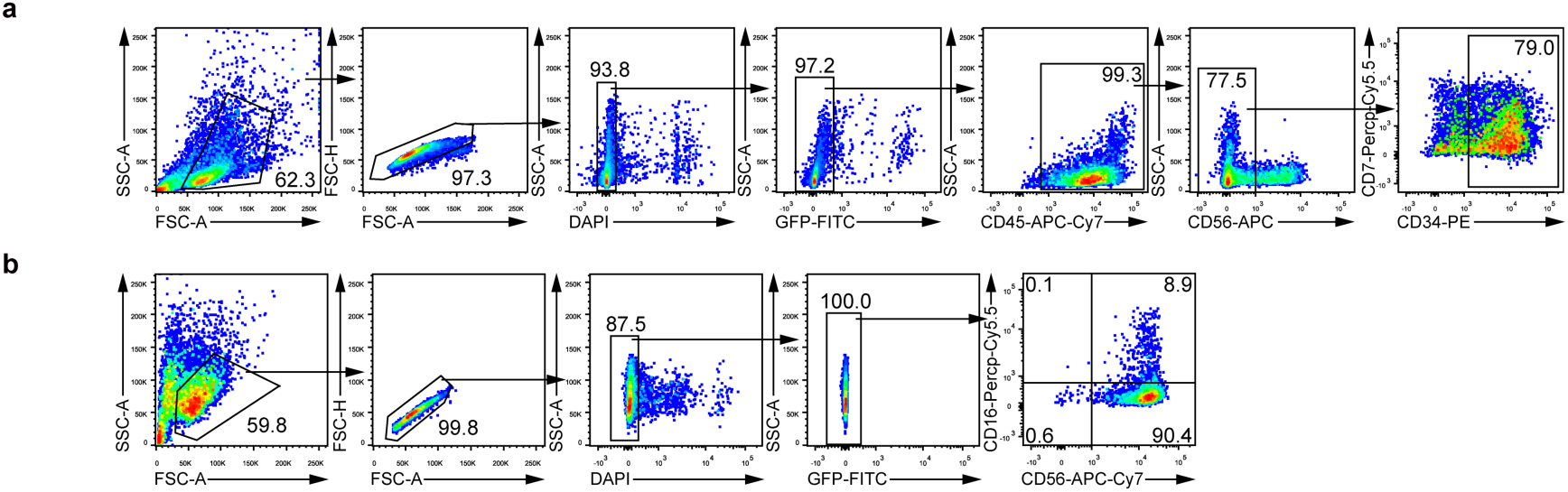
Gating strategy of day-19 iNKP and derived iNK from 14-day culture *in vitro*. **a,** Gating strategy of iNKP cells from day-19 organoids. **b,** Gating strategy of iNK cells matured from day-19 iNKP *in vitro*.

**Extended Data Fig. 3.**
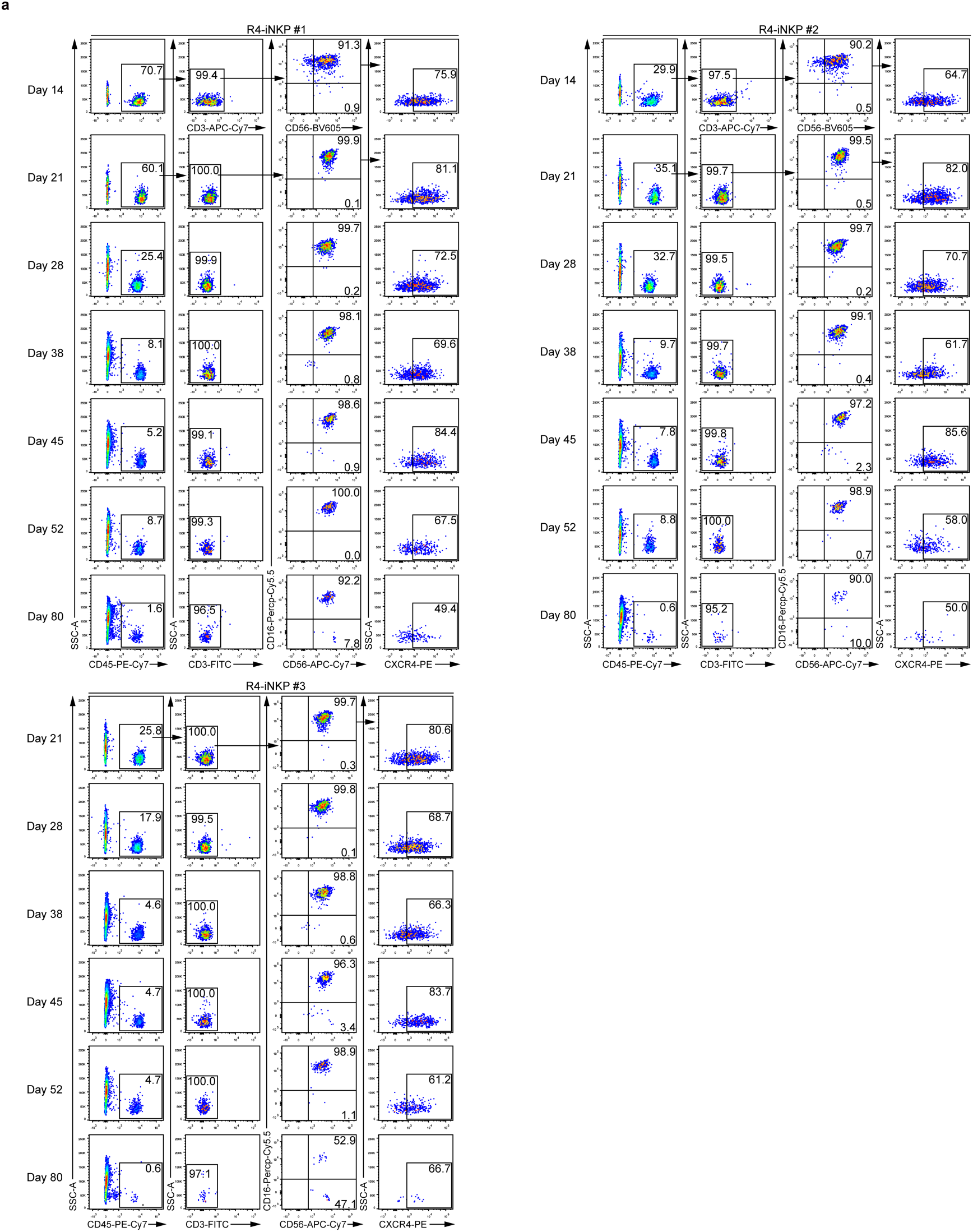
Flow cytometry of iNK in PB of R4-iNKP cell recipients. **a,** Flow cytometry of iNK cells in PB from the R4-iNKP cell recipients (n = 3 animals) on days 14, 21, 28, 38, 45, 52, and 80 post-infusions.

**Extended Data Fig. 4.**
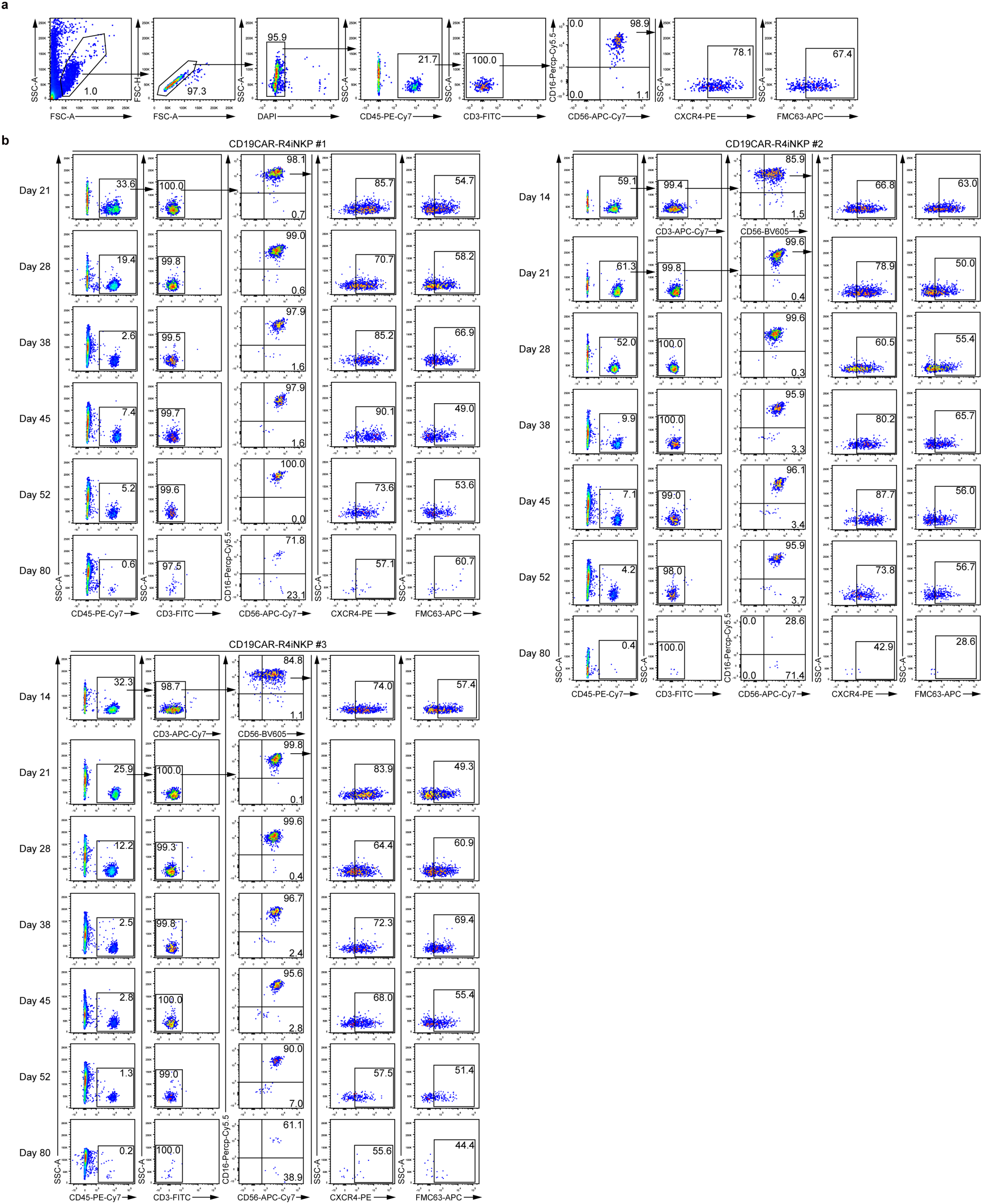
Flow cytometry of iNK cells in PB of CD19CAR-R4iNKP cell recipients. **a,** Gating strategy of iNK cells in PB from a representative CD19CAR-R4iNKP cell recipient. **b,** Flow cytometry of iNK cells in PB from CD19CAR-R4iNKP cell recipients (n = 3 animals).

**Extended Data Fig. 5.**
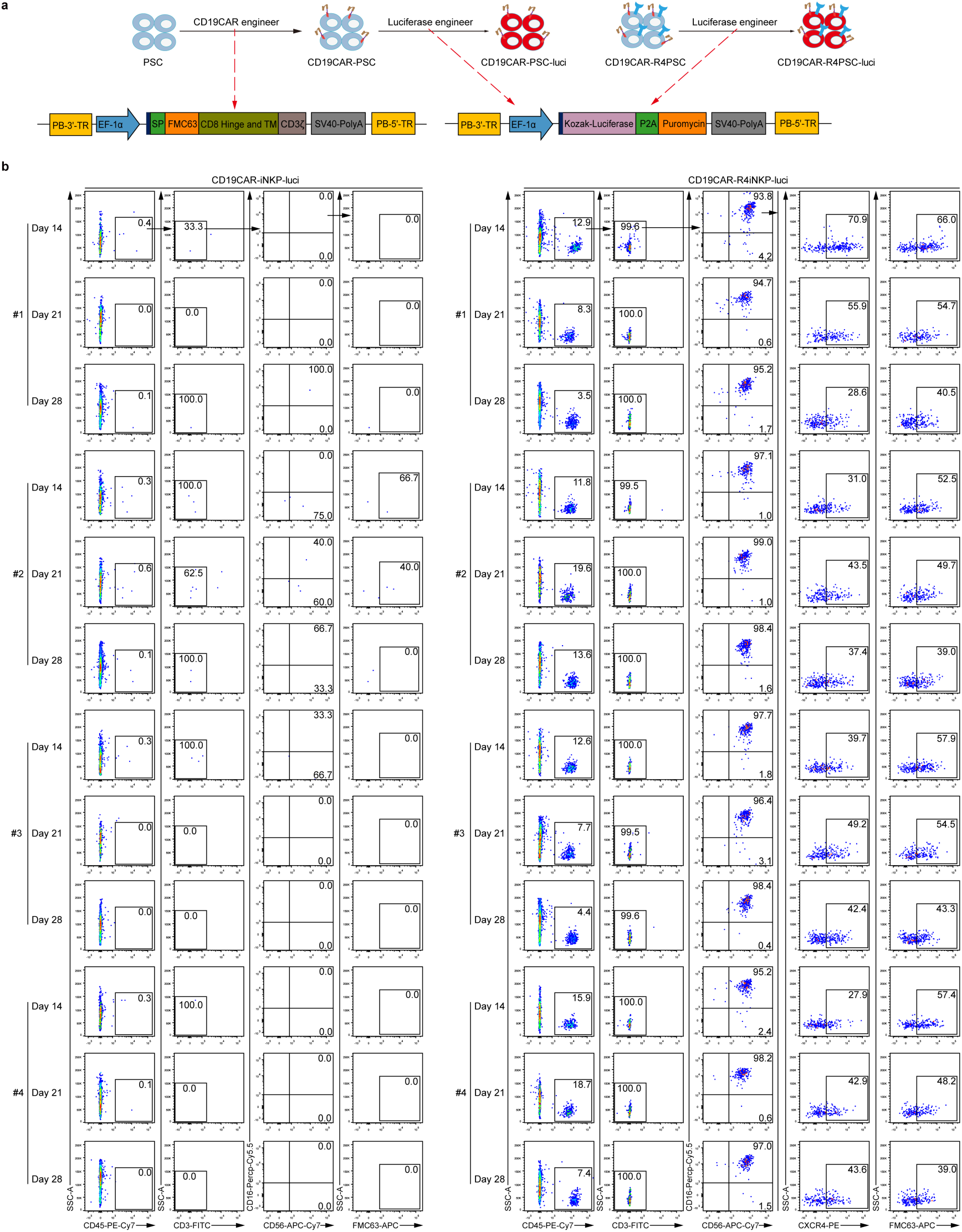
Flow cytometry of iNK cells from the CD19CAR-iNKP-luci cell recipients and CD19CAR-R4iNKP-luci cell recipients. **a,** Schematic diagram of generating luciferase-expressing CD19CAR-PSC (CD19CAR-PSC-luci) line and luciferase-expressing CD19CAR-R4PSC (CD19CAR-R4PSC-luci) line. **b,** Flow cytometry of the iNK cells in PB from CD19CAR-iNKP-luci cell recipients (n = 4 animals) and CD19CAR-R4iNKP-luci cell recipients (n = 4 animals).

**Extended Data Fig. 6.**
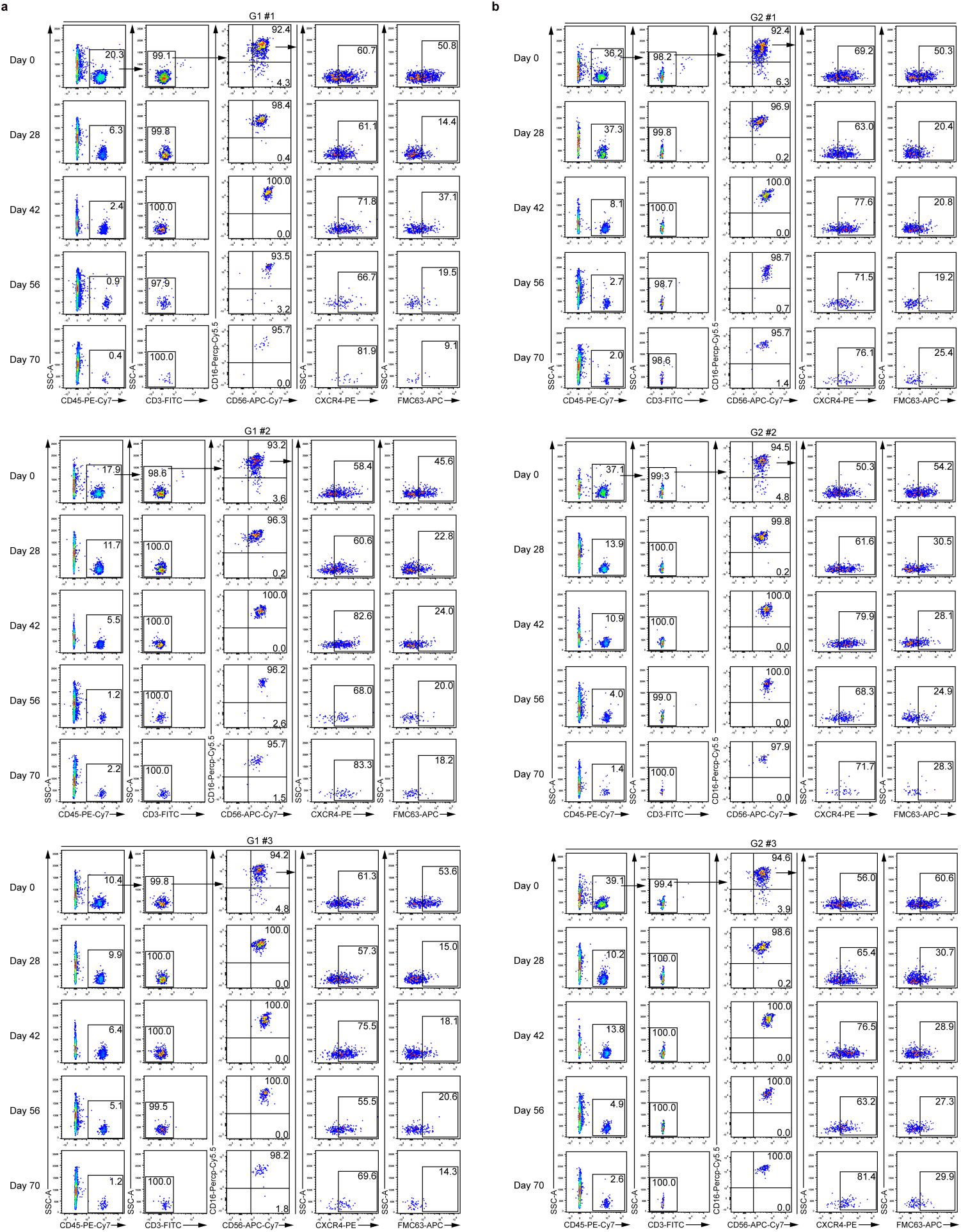
Flow cytometry of iNK cells from CD19CAR-R4iNKP cell recipients. **a,** Flow cytometry of iNK in PB of CD19CAR-R4iNKP cell recipients (n = 3 animals) from G1 on days 0, 28, 42, 56, and 70 after Nalm-6-luci cell injection. **b,** Flow cytometry of iNK in PB of CD19CAR-R4iNKP cell recipients (n = 3 animals) from G2 on days 0, 28, 42, 56, and 70 after Nalm-6-luci cell injection.

**Extended Data Fig. 7.**
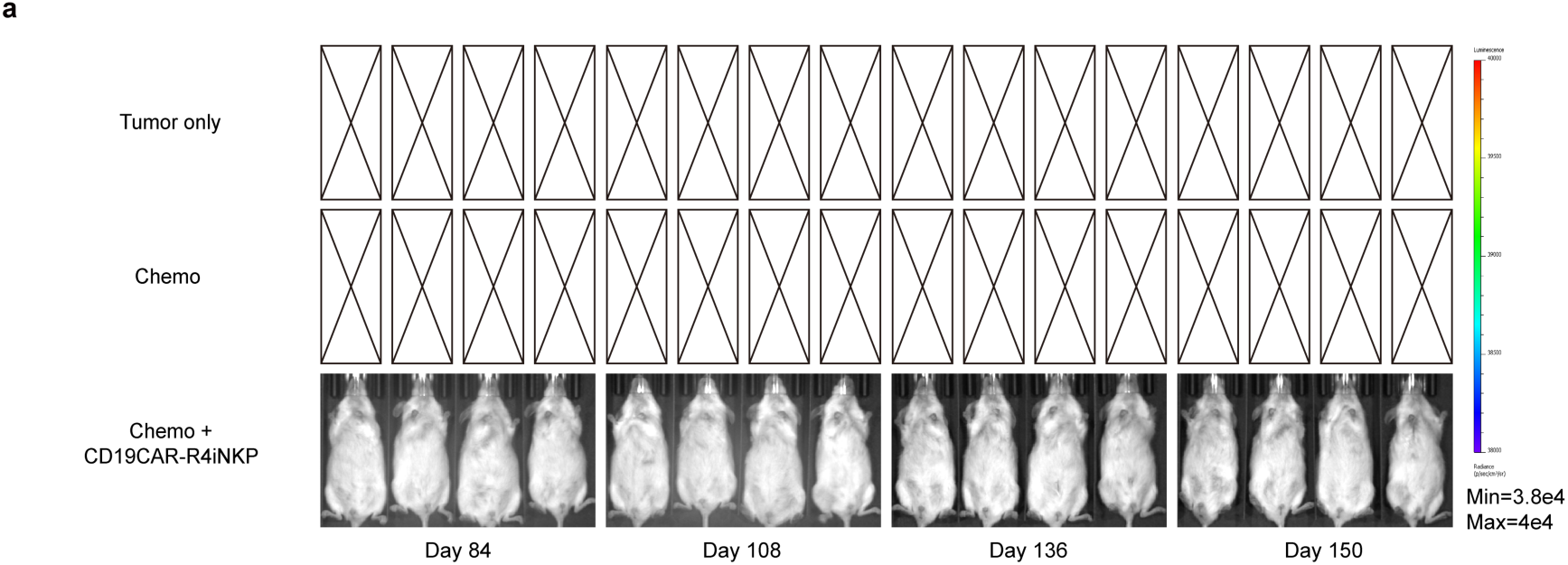
Prolonged monitoring of the kinetics of tumour burden of tumour-bearing mice treated with Chemo and CD19CAR-R4iNKP. **a,** BLI of the tumour-bearing mice from Chemo + CD19CAR-R4iNKP group (n = 4 animals) on days 84, 108, 136, and 150.

**Extended Data Fig. 8.**
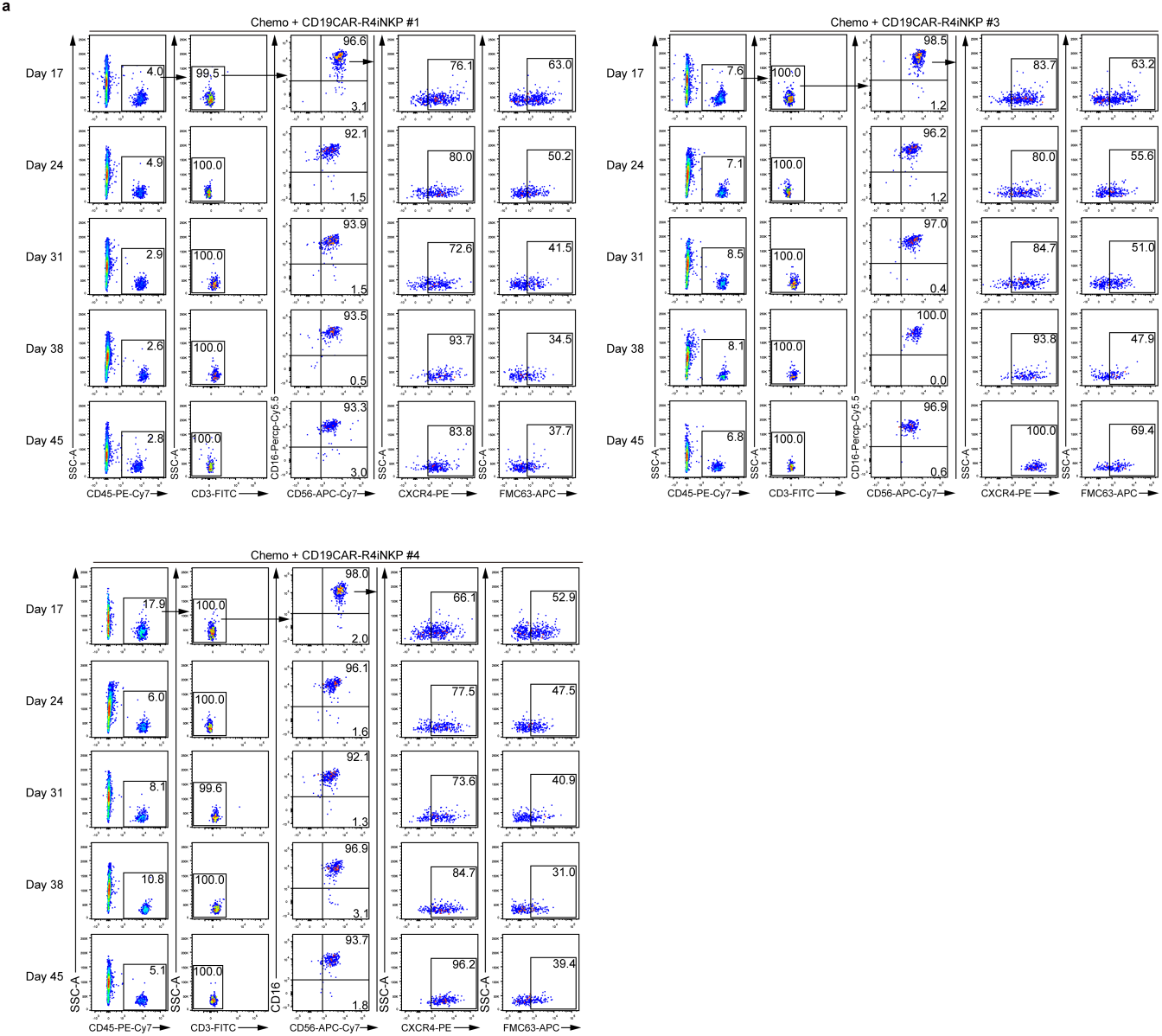
Flow cytometry of iNK cells in PB of the tumour-bearing mice after Chemo and CD19CAR-R4iNKP combination therapy. **a,** Flow cytometry of iNK cells in PB from tumour-bearing mice (n = 4 animals) after Chemo and CD19CAR-R4iNKP combination therapy on days 17, 24, 31, 38, and 45.

## Notes

### Competing Interest Statement

The authors have declared no competing interest.

